# Decoration of *Burkholderia* Hcp1 protein to virus-like particles as a vaccine delivery platform

**DOI:** 10.1101/2024.01.17.576127

**Authors:** Nittaya Khakhum, Noe Baruch-Torres, Jacob L. Stockton, Itziar Chapartegui-González, Alexander J Badten, Awadalkareem Adam, Tian Wang, Alejandro Huerta-Saquero, Y. Whitney Yin, Alfredo G Torres

## Abstract

Virus-like particles (VLPs) are protein-based nanoparticles frequently used as carrier in conjugate vaccine platforms. VLPs have been used to display foreign antigens for vaccination and to deliver immunotherapeutic against diseases. Hemolysin-coregulated proteins 1 (Hcp1) is a protein component of the *Burkholderia* type 6 secretion system which participates in intracellular invasion and dissemination. This protein has been reported as a protective antigen and is used in multiple vaccine candidates with various platforms against melioidosis, a severe infectious disease caused by the intracellular pathogen *B. pseudomallei*. In this study, we used P22 VLPs as a surface platform for decoration with Hcp1 using chemical conjugation. C57BL/6 mice were intranasally immunized with three doses of either PBS, VLPs or conjugated Hcp1-VLPs. Immunization with Hcp1-VLPs formulation induced Hcp1-specific-IgG, IgG_1_, IgG_2c_ and IgA antibody responses. Furthermore, the serum from Hcp1-VLPs immunized mice enhanced the bacterial uptake and opsonophagocytosis by macrophages in the presence of complement. This study demonstrated an alternative strategy to develop a VLPs-based vaccine platform against *Burkholderia* species.

## Introduction

Viral like particles (VLPs) are non-infectious nanocage architectures and self-assemble particles which express genome-encoded protein capsids, core and the viral envelop but cannot replicate because the genetic material is lacking (1). VLPs are broadly used in medical biotechnology applications including drug delivery, vaccine platforms, gene regulation, gene delivery and imaging (2). For example, the *Salmonella enterica* serovar Typhimurium bacteriophage P22 VLP is approximately 60 nm in size with and icosahedral shape assembled with 420 copies of the 46 kDa coat protein (CP) and 300 copies of the 34 kDa scaffolding protein (SP) (3).

The encapsulation of P22 VLP with proteins of interest has been developed as nanoreactors (or nanocages) and used for drug and vaccine delivery platform against diseases (3-7). The strategy of this method is the fusion of the cloned gene encoding the protein of interest to SP gene followed by recombinant expression together with CP and *in vivo* assembly to generate a nanoreactor (4). However, this methodology is mostly suitable for soluble protein of interest especially in the protein purification step. For the insoluble protein, it is more difficult and challenging to purify nanoreactors without aggregation. An external decoration of VLPs with target protein is an alternative method by using two classic chimerical vaccine design approaches of genetic insertion and chemical conjugation (8). Genetic insertion is easily achieved by genetic engineering and antigens are presented on the VLPs surface with high efficiency, but sometimes they form inclusion bodies and takes longer for the production process (9). For the chemical conjugation, covalently attached antigens on the VLPs surface have higher efficiency to induce immune response compared to non-covalent conjugation (10).

Two features, size (20 – 200 nm), and repetitive surface geometrical structure of VLPs are critical for their ability to potently activate B cells and elicit strong and long-lasting antibody responses which are important to their advantages as a vaccine (11, 12). VLPs enter lymphatic vessels and passive drainage to the subcapsular region of lymph nodes and optimal uptake by antigen presenting cells (APCs) (13), following the T helper cells activation and antigen presentation on MHC class II molecules (14). Moreover, trafficking of VLPs to B cell follicles is facilitated through specific interactions between VLPs and complement components or natural IgM antibodies (15). VLPs are highly stable with variation of temperature and other conditions. However, VLPs have been modified to enhance structural stability using engineering techniques such as intersubunit disulfide bonds for thermostability purpose (16-18). The P22 VLP has been reported to elicit a strong protective immune response in mice against influenza viruses without the addition of adjuvants (19).

The hemolysin-coregulated proteins (Hcp) are components of Gram-negative bacterial type 6 secretion systems (T6SS) (20). This protein forms hexameric rings and cooperates with other proteins in the T6SS to puncture neighboring host cells to deliver a cargo molecule. Hcp1 protein of *Burkholderia pseudomallei*, an intracellular Gram-negative bacterium that causes melioidosis disease, has been reported as a protective antigen (21). Interestingly, this protein has been used to construct various vaccine platforms against pathogenic *Burkholderia* with different outcomes depending on the platform, including gold-nanoparticles (22-24), glycoconjugate subunits (25) and membrane vesicles (26).

In this study, we develop an approach for exterior decoration of the Hcp1 protein on the P22 VLPs surface by the chemical conjugation method and evaluate the antibody induction following immunization of mice. The conjugation was analyzed under transmission electron microscope (TEM) before immunization. Protein specific antibody titer was quantified and the ability of immune serum to promote opsonophagocytic activity by macrophages was further evaluated.

## Materials and methods

### Hcp1 expression and purification

*Burkholderia*-specific protein Hcp1 (BMAA0742) construction and expression was described elsewhere (22, 24) with modifications. The *hcp1* gene was cloned into the pET30a(+) containing a kanamycin resistance gene. Protein expression was conducted in an *E. coli* BL21(DE3) by growing an overnight preculture in LB media containing 50 μg/ml kanamycin. Then a larger LB culture was inoculated with the preculture at a dilution of 1:100 and incubated at 37ºC, 200 rpm (LabLine 3525). The culture was induced with 1 mM of isopropyl-β-D-1-thiogalactopyranoside (IPTG) after an optical density at 600 nm (OD_600_) reached 0.6. After 4 h induction at 37ºC, the culture was centrifuged, and each resulting bacterial pellet was frozen at −80°C. The bacterial pellets were then resuspended in 20 ml of lysis buffer pH 8.0 (50 mM NaH_2_PO_4_, 300 mM NaCl, 5 mM Imidazole) containing 1× cOmplete EDTA-free protease inhibitor cocktail (Roche, Germany) and 1 mg/ml lysozyme. The lysate was then sonicated and centrifuged (10,000 ×g, at 4°C for 30 min), and the supernatant was filtered (0.2-μm pore size; Millipore). Soluble protein fractions were then mixed with pre-equilibrated Ni-NTA agarose resin (Qiagen) and incubated at 4°C for 1 h with rotation. The incubated resin was loaded into 5 ml polypropylene column (Qiagen) subsequently multiple washes were applied using a wash buffer pH 8.0 (50 mM NaH_2_PO_4_, 300 mM NaCl, 10 mM Imidazole and 1× cOmplete EDTA-free protease inhibitor cocktail) and eluted with an elution buffer pH 8.0 (50 mM NaH_2_PO_4_, 300 mM NaCl, 150 mM Imidazole). Fractions containing soluble protein were pooled before dialysis into PBS overnight at 4°C. The protein was concentrated using a 10 kDa MWCO Amicon concentrator device (Millipore). The purified proteins and protein standards were subjected to a colorimetric bicinchoninic acid (BCA) assay following the manufacturer’s protocol and then stored at −80°C until use (Pierce protein assay kit; ThermoFisher Scientific, USA). The proteins were visualized by running a 4-20% SDS-PAGE (Bio-Rad) and stained with Coomassie blue.

### P22 VLPs production and purification

*E. coli* BL21 (DE3) carrying pBAD-P22-SP and pRSF/P22-CP plasmids was grown overnight in LB medium then diluted to 1:100 in terrific broth (TB) medium before incubated at 37°C, 200 rpm (LabLine 3525) until OD_600_ reached 0.6. The SP-protein expression was induced with 0.125% L-arabinose for 16 h at 27°C, 120 rpm (LabLine 3525), and subsequently, the CP was induced with 0.5 mM IPTG for 4 h at 37ºC. The bacterial pellet was collected by centrifugation then resuspended in PBS (pH 7.4), and sonicated. The resulting suspension was centrifuged at 12,000 ×g for 30 min at 4°C and the clear lysate was collected before ultracentrifuged through 35% w/v sucrose in PBS at 274,000 ×g, 4°C for 1 h in a SW 41 Ti rotor (Beckman Coulter, Inc). The pellet was resuspended in 1 ml of PBS pH 7.4, centrifuge at 9,300 ×g and further purified using a Sephacryl S-500, 60 × 1.6 cm size exclusion column (GE Healthcare) in FPLC (AKTA Pharmacia). Collected fractions containing well-formed VLPs were analyzed by SDS-PAGE and under TEM.

### Chemical functionalization of P22 VLPs with Hcp1 protein

Hcp1 proteins were covalent conjugated to P22 VLPs by the carbodiimide crosslinker (18). 53.36 nmoles of the purified Hcp1 (18.74 kDa) was mixed with 5,336.00 nmoles of (3-Dimethylaminopropyl)-N′-ethylcarbodiimide hydrochloride (EDC) (Sigma) and 5,336.00 nmoles N-hydroxysuccinimide (NHS) (Sigma) (Protein: EDC or NHS ratio is 1:100) in PBS pH 5.5, then the mixture was slowly rotated at room temperature (RT) for 30 min. The mixture was filtered through a 10 kDa Amicon concentrator (Millipore) to remove the excess of EDC and NHS. The filtered Hcp1 was mixed with 21.27 nmoles P22 VLPs (47 kDa) at a ratio of 1:10 followed by adding each EDC and NHS (VLPs: EDC or NHS ratio is 1:600) in PBS pH 7.4 (Fig. 1A). The reaction was kept in slow rotation at RT for 3 h. Five milliliters of PBS pH 7.4 were added into the reaction mixture to scale up volume before loading into centrifuge tubes containing 35% sucrose solution. The final Hcp1-VLPs particles were pelletized by ultracentrifugation at 274,000 ×g at 4°C for 1 h in a SW41 Ti rotor (Beckman Coulter, Inc). The final pellet was resuspended in PBS pH 7.4.

**Fig. 1.**
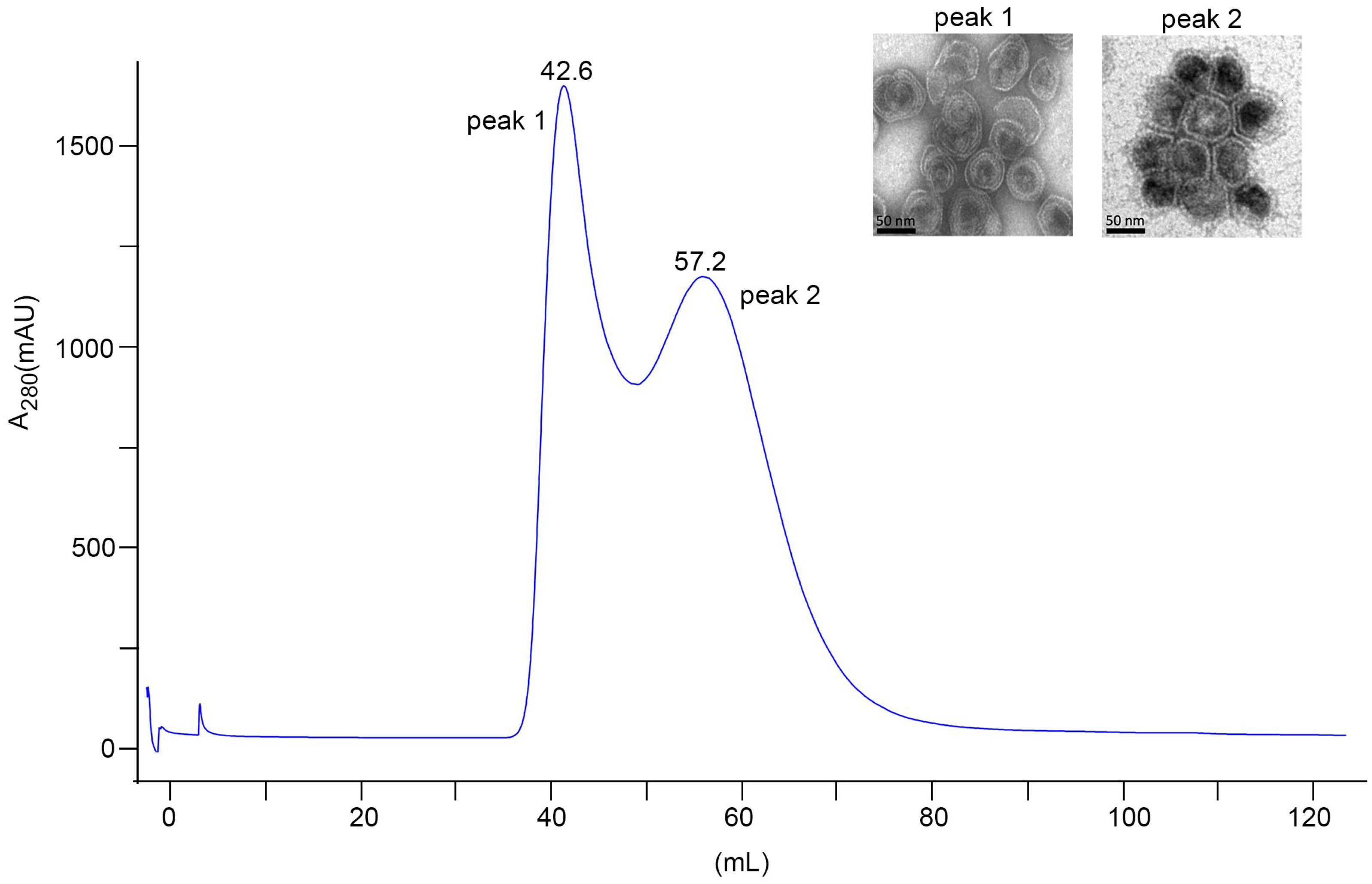
Purification of P22 VLPs. Size exclusion chromatography profile of P22 VLPs purification on a HiPrep 16/60 Sephacryl S-500HR. Sample was loaded followed by 1 Column volume of PBS. Two milliliters of eluted fractions were collected. Peak 2 (42.6 mL) shows fraction that contains incomplete assembled VLPs or larger aggregated VLPs and the peak 2 (57.2 mL) shows well-formed VLPs. Representative fractions from the peak 1 and peak 2 were applied onto copper grid and stained with 2% aqueous uranyl acetate. The morphology was analyzed under TEM. The scale bars represent 50 nm.

### Negative staining and Immunogold labeling TEM

Unconjugated VLPs and Hcp1-VLPs conjugated suspension (10 µl) were loaded onto CF200-CU carbon film 200 mesh copper grid (Electron Microscope Science) and incubated at RT for 10 min. For negative staining, grids were blotted with filter paper before being placed onto 2% aqueous uranyl acetate for 1 min, blotted and dried under the light. For immunogold labeling, grids were placed on the surface of 1:1000 primary antibody (Mouse monoclonal [HIS.H8] antibody, Abcam) and incubated for 30 min at RT then washed with 1% BSA (w/v) in Tris-buffered saline (TBS). The grids were stained with 1:20 secondary antibody (6 nm immunogold goat-anti-Mouse IgG (H&L), Electron Microscopy Sciences Aurion) for 30 min at RT. After incubation, grids were washed and fixed with 2% glutaraldehyde for 10 min, then dried before imaging under TEM.

### Intranasal immunization and blood collection

Female 6-8-week-old C57BL/6 mice (n = 5/group) were intranasally immunized with 3 doses (prime and 2 boosts) of PBS, unconjugated VLPs or Hcp1 conjugated VLPs (5 µg or 10 µg). Blood from individual mouse were collected via retro-orbital (RO) bleeding a week before prime (baseline) and 2 weeks after the 2^nd^ boost (post). Blood was stored in Microvette tubes without an anticoagulant (SARSTEDT) and incubated at room temperature for 30 min to permit clotting before centrifugation. The serum was collected and stored at -80°C until further analysis.

### Detection of Hcp1-specific antibodies

The Hcp1-specific IgG, IgG_1_, IgG_2c_ and IgA endpoint titers were detected using indirect ELISA. Briefly high binding 96-well plates were coated with 1 µg/well of purified Hcp1 protein at 4°C overnight. Coated plates were washed (0.05% Tween 20 in 1× DPBS) and incubated with blocking buffer (8% heat inactivated (HI) fetal bovine serum (FBS) in 1× DPBS) at RT for 2 h. After blocking, plates were washed and incubated with two-fold dilution of baseline or post-sera collected from immunized mice at RT for 2 h. Plates were washed before incubation with 1:5,000 dilution of goat anti-mouse IgG, IgG_1_, IgG_2c_ or IgA secondary antibody (Southern Biotech) at RT for 2 h. Tetramethylbenzidine (TMB) substrate solution (Invitrogen) was added and incubated for 10 min before adding the stop solution (2 N H_2_SO_4_). Plates were read at 450 and 570 nm using a microplate reader (Biotek). The results were reported as the reciprocal of the highest titer giving an optical density (OD) reading of at least mean 2 SD from the mean baseline sera OD. All assays were performed in triplicate.

### Macrophage infection and opsonophagocytosis

RAW 264.7 (ATCC TIB-71) macrophages were routinely grown in complete DMEM (cDMEM) (Gibco) supplemented with 10% HI FBS, sodium pyruvate, non-essential amino acid, and antibiotics (penicillin and streptomycin) at 37°C under an atmosphere of 5% CO_2_. The day prior to cell infection, RAW 264.7 cells were seeded at a density of 5 × 10^5^ cells/well in 24-well tissue culture plates (ThermoFisher) for bacterial uptake assay and in 12-well plates with round coverslips for immunofluorescent microscopic analysis. The following day, a 12 h overnight culture of *B. pseudomallei* K96243 was diluted to 5 × 10^6^ CFU with cDMEM without antibiotic equivalent to a multiplicity of infection (MOI) of 10. The bacteria were incubated with PBS (no sera) or 10% of sera with and without HI that were pooled from immunized mice (PBS, unconjugated VLPs, 5 µg or 10 µg of Hcp1 conjugated VLPs) at 37°C for 1 h with slightly agitation. The opsonized bacterial suspension was added to pre-washed RAW264.7 monolayers prior to incubation at 37°C, 5% CO_2_ for 2 h. For bacterial uptake enumeration, the monolayers were washed twice with PBS to remove extracellular bacteria then lysed with 0.2% Triton X-100 (Sigma) in PBS before serial dilution and plating on LB agar. Each experimental group was assayed in triplicate, and three independent experiments were performed.

### Immunofluorescent staining and microscopic analysis

Infected RAW 264.7 cells were fixed with 4% paraformaldehyde–PBS for 30 min following the select agent inactivation protocol approved by UTMB Biosafety committee. Cells were permeabilized with 0.25% Triton X-100 in PBS for 7 min at RT before incubation with serum (1:1000) from *B. pseudomallei* PBK001 live attenuated vaccine immunized mice (27). Cells were washed then incubated with 1:5,000 goat anti-mouse IgG, IgM (H+L) secondary antibody conjugated to Alexa 488 (Invitrogen) followed by actin and DNA staining using rhodamine phalloidin (Invitrogen) and DAPI (Sigma) together at 1:10,000 dilutions. The coverslips were mounted using Prolong gold antifade (Molecular Probes, Life Technology) and sealed with fixative. Stained cells were visualized using an Olympus BX51 upright fluorescence microscope and analyzed using ImageJ software from the National Institutes of Health (NIH).

## Results

### P22 VLPs production and purification

The P22 VLPs were produced by two steps protein induction. The pBAD-P22-SP plasmid was first induced to obtain P22 scaffold protein followed by induction of pRSF/P22-CP plasmid for capsid protein. The P22 VLPs were purified from the bacterial suspension using 35% sucrose cushion ultracentrifugation followed by gel filtration chromatography. Two protein-containing fractions were eluted after running through Sephacryl S-500 size exclusion column. Each of these fractions was morphologically characterized via TEM. The TEM imaging indicates that the first fraction contains incompletely assembled VLPs while the second fraction contains well-formed VLPs (Fig. 1). Peak 1 shows VLPs that are not in the right conformation due to protein aggregations that make these VLPs close to the void volume, which indicates a high number of defective subunits. The aggregation occurs during spontaneously assembly of VLPs inside the expression host and they may contain desired compound and disassembled form of VLPs. While the elution fraction from the peak 2, corresponding to the well-formed VLPs, contains a defined number of subunits folded properly that can be eluted at a different retention volume with a small molecular size. Therefore, the elution fractions from peak 2 containing highly pure assembled VLPs which were subsequently used for the protein conjugation step.

### Chemical functionalization of P22 VLPs with Hcp1 protein

The free carboxyl group of Hcp1 was activated with carbodiimide and N-hydroxysuccinimide in acidic conditions (pH 5.5) to prevent the self-conjugation of Hcp1 homodimers and to prevent hydrolysis of the activated functional groups. Hcp1 was subsequently coupled to the free amine groups exposed on P22 VLPs (Fig. 2A). To evaluate the efficiency of conjugation, the conjugated number of conjugated Hcp1 molecules on the surface of the VLPs was quantified using TEM (with 20 fields) as shown in Table S1. The average total conjugation efficiency was about 24%. The purified Hcp1 (19.54 kDa), VLPs capsid protein (47 kDa), before and after chemical conjugation in different concentration, were visualized on an SDS-PAGE gel followed by Coomassie blue staining (Fig. S1). Negative staining on TEM showed homogeneous hexahedral structures of VLPs with an average diameter of 50 nm (Fig. 2B). To evaluate the successful Hcp1 conjugation to the surface of VLPs, we probed the recombinant Hcp1 with anti-His tag primary antibody followed by 6 nm gold nanoparticle conjugated goat-anti-Mouse IgG secondary antibody and the surface exposed proteins were labeled (Fig. 2C).

**Fig. 2.**
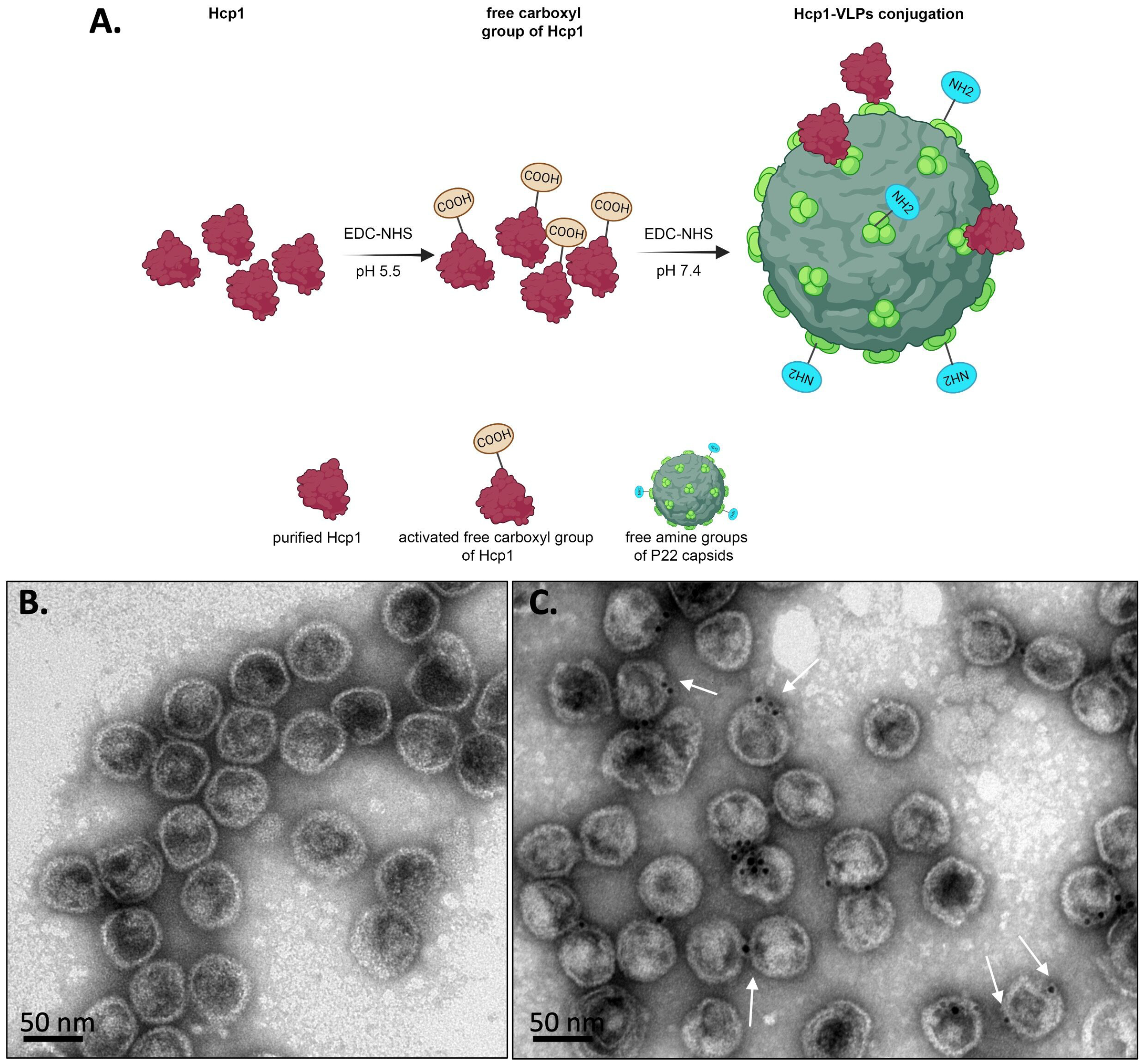
Schematic showing the methodology used to conjugate Hcp1 protein to P22 VLPs. (**A**) Two steps of chemical functionalization of P22 VLPs with Hcp1. Free carboxy groups of Hcp1 protein and VLPs are released after exposing to EDC and NHS chemistry at pH 5.5 and 7.4, respectively followed by the covalent conjugation of Hcp1-VLPs. (**B**) Negative staining and (**C**) immunogold labeling of Hcp1-VLPs conjugation under TEM. The white arrows indicate the Hcp1 molecule covalently attached to the VLP surface. The scale bars represent 50 nm. The figure 2A was created with Biorender (biorender.com).

### Intranasal immunization of Hcp1-VLPs conjugation induces protein specific antibodies response

The timeline immunization protocol is shown in Fig. 3. The endpoint titers of Hcp1 specific serum IgG, IgG_1_, IgG_2c_ and IgA were assessed by ELISA using purified Hcp1 as the coating protein antigen. The ELISA results demonstrated that mice immunized with low (5 µg) and high (10 µg) dose of Hcp1-VLPs conjugation stimulated the production of high titers of IgG (∼10^6^), IgG_1_ (∼10^5^-10^6^), IgG_2c_ (∼10^5^) subclass and IgA (∼10^4^) without an apparent difference between doses (Fig. 4A–D). In contrast, serum from mice receiving PBS and VLPs showed low titers (50 - 100) in all detected antibodies.

**Fig. 3.**
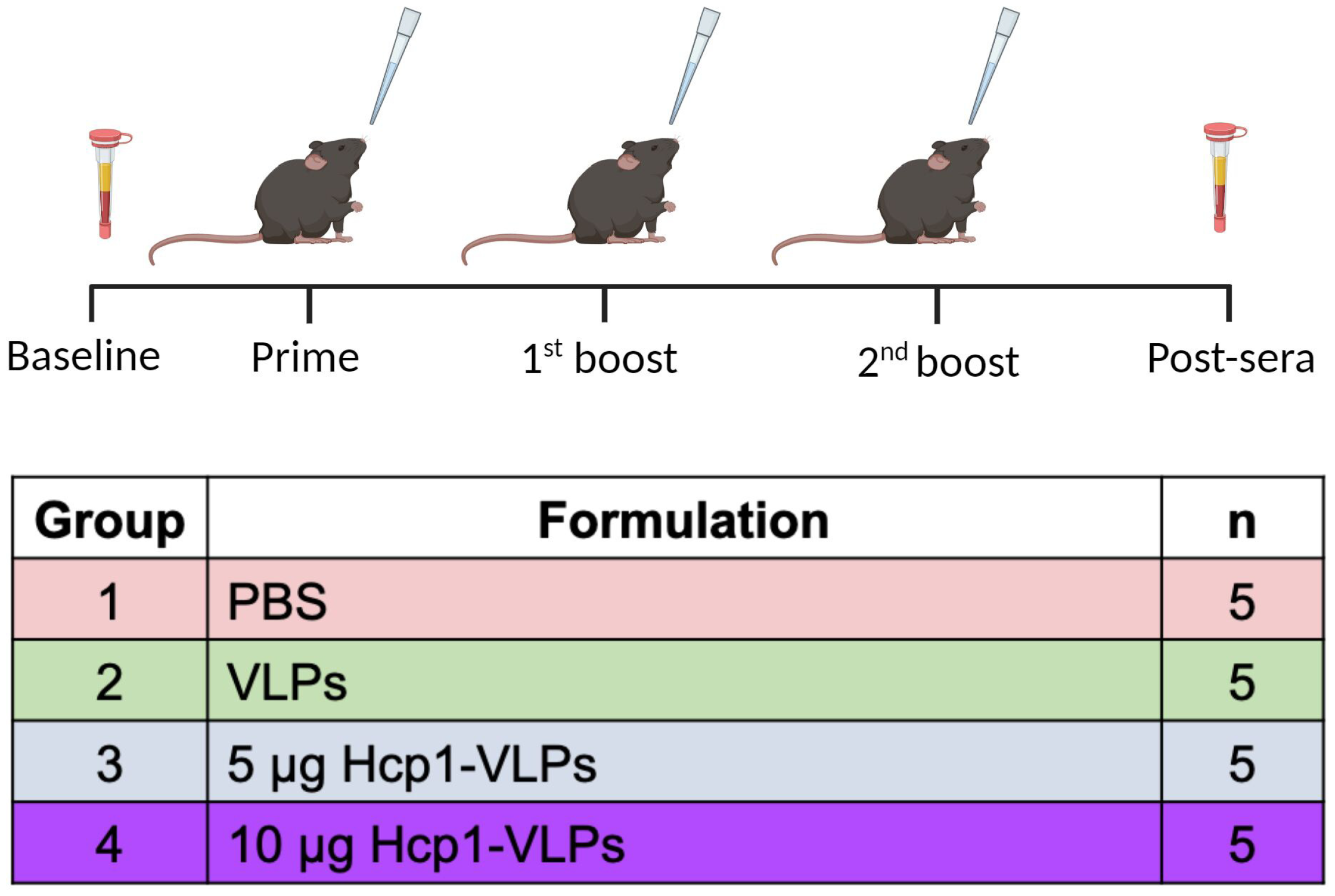
Timeline of immunization and sample collection. Female C57BL/6 mice (n = 5/group) received prime and boosts with PBS, VLPs, 5 or 10 µg Hcp1-VLPs formulation, via the intranasal route, at two-week intervals. Baseline and post-immunization serum were collected a week prior to prime and two weeks after the 2^nd^ boost, respectively. The figure was created with Biorender (biorender.com).

**Fig. 4.**
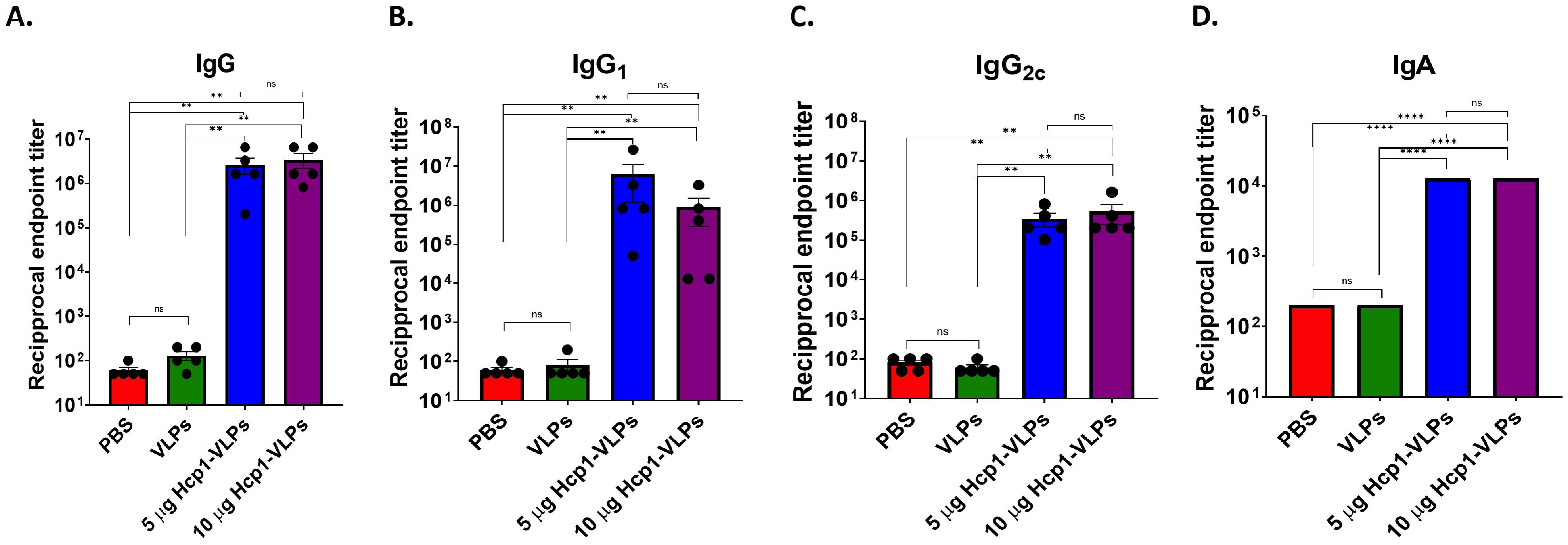
Intranasal immunization of Hcp1-VLPs conjugation induces protein specific antibodies response. Mice were intranasally immunized with PBS, VLPs, 5 or 10 µg Hcp1-VLPs. Baseline and post-immunization sera were collected and used for analysis of Hcp1 specific IgG (**A**), subclass IgG_1_ (**B**), IgG_2c_ (**C**) and IgA (**D**) by indirect ELISA. The results were reported as the reciprocal of the highest titer giving an optical density (OD) reading of at least 2 SD from the mean baseline sera OD. All assays were performed in triplicate. The limit of detection (LOD) is 50. Data are analyzed using Mann-Whitney test. Bars represent means with standard error of the mean (SEM). Values that are significantly different are indicated by bars and asterisks as follows: **, *P* < 0.01, ****, *P* < 0.0001, n.s., not significant.

### Serum from Hcp1-VLPs immunized mice enhances opsonized bacterial uptake by macrophages in the presence of complement

To assess the functionality of serum antibodies, we evaluated whether the antibodies promote antibody-mediated bacterial opsonophagocytosis of *B. pseudomallei*. Pooled serum from different immunization groups were pre-incubated with *B. pseudomallei* prior to infect RAW

246.7 macrophages. After infection, percent bacterial uptake in macrophages was quantified by standard dilution and plating method. The results demonstrated that the heat-inactivated (HI) serum from PBS, VLPs, 5 and 10 µg of Hcp1-VLPs group had no significant effect on percent of the bacterial uptake among groups as well as no sera control group (Fig. 5). In contrast, the presence of complement in the no-HI serum of both doses of Hcp1-VLPs group increased the bacterial uptake by macrophages whereas the serum from PBS and VLPs group showed no significant effect compared to those two groups (Fig. 5). Moreover, the results indicated the high dose (10 µg) of Hcp1-VLPs had a higher ability to enhance bacterial uptake compared to low dose (5 µg) (Fig. 5). To confirm the bacterial quantification results, the infected cells were fixed, permeabilized, stained then visualized using immunofluorescence microscopy. A higher number of *B. pseudomallei* were accumulated and internalized by macrophages in the presence of no-HI serum from animal immunized with 5 and 10 µg of Hcp1-VLPs groups than others which had less bacteria (Fig. 6). These immunofluorescent images confirmed the bacterial uptake results and demonstrated that the Hcp1-VLPs formulation induced antibody response in promoting opsonophagocytosis that was mediated by complement.

**Fig. 5.**
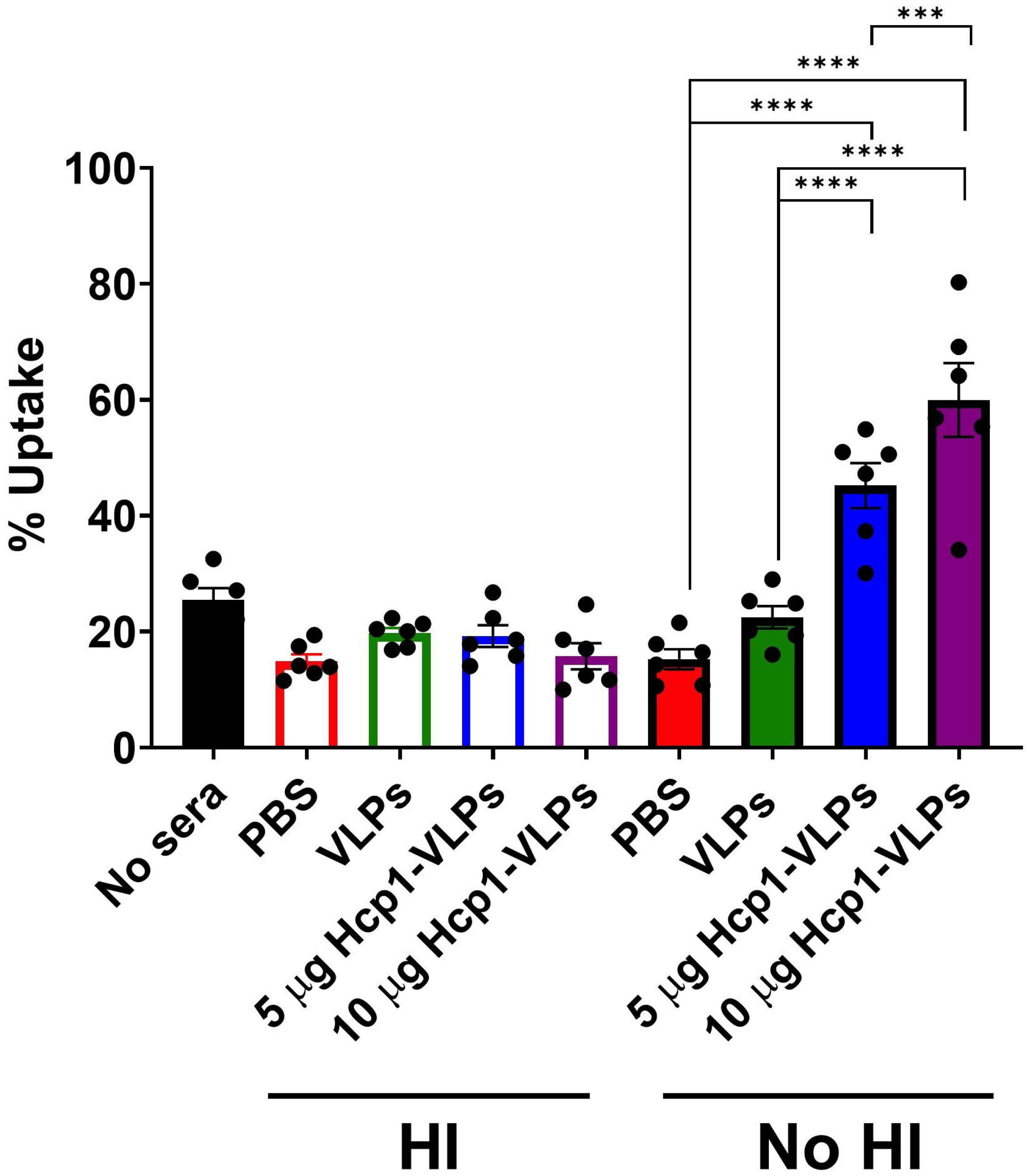
Serum from Hcp1-VLPs immunized mice enhance the uptake of opsonized bacteria by macrophages in the presence of complement. Opsonophagocytosis was performed by incubating cDMEM (no sera), heat-inactivated (HI) or no-HI pooled immune sera from mice (n = 5) receiving PBS, VLPs, 5 or 10 µg Hcp1-VLPs and incubating with *B. pseudomallei* K96243 for 1 h. After incubation, opsonized bacteria were used to infect RAW 264.7 macrophage cell monolayers at 37°C with 5% CO_2_ for 2 h. After infection, the bacterial uptake by macrophages was enumerated. The values of opsonophagocytosis assay are represented as mean ± SEM from two individual assays conducted in triplicate. Significant differences were determined using a one-way ANOVA. ***, *P* < 0.001, ****, *P* < 0.0001.

**Fig. 6.**
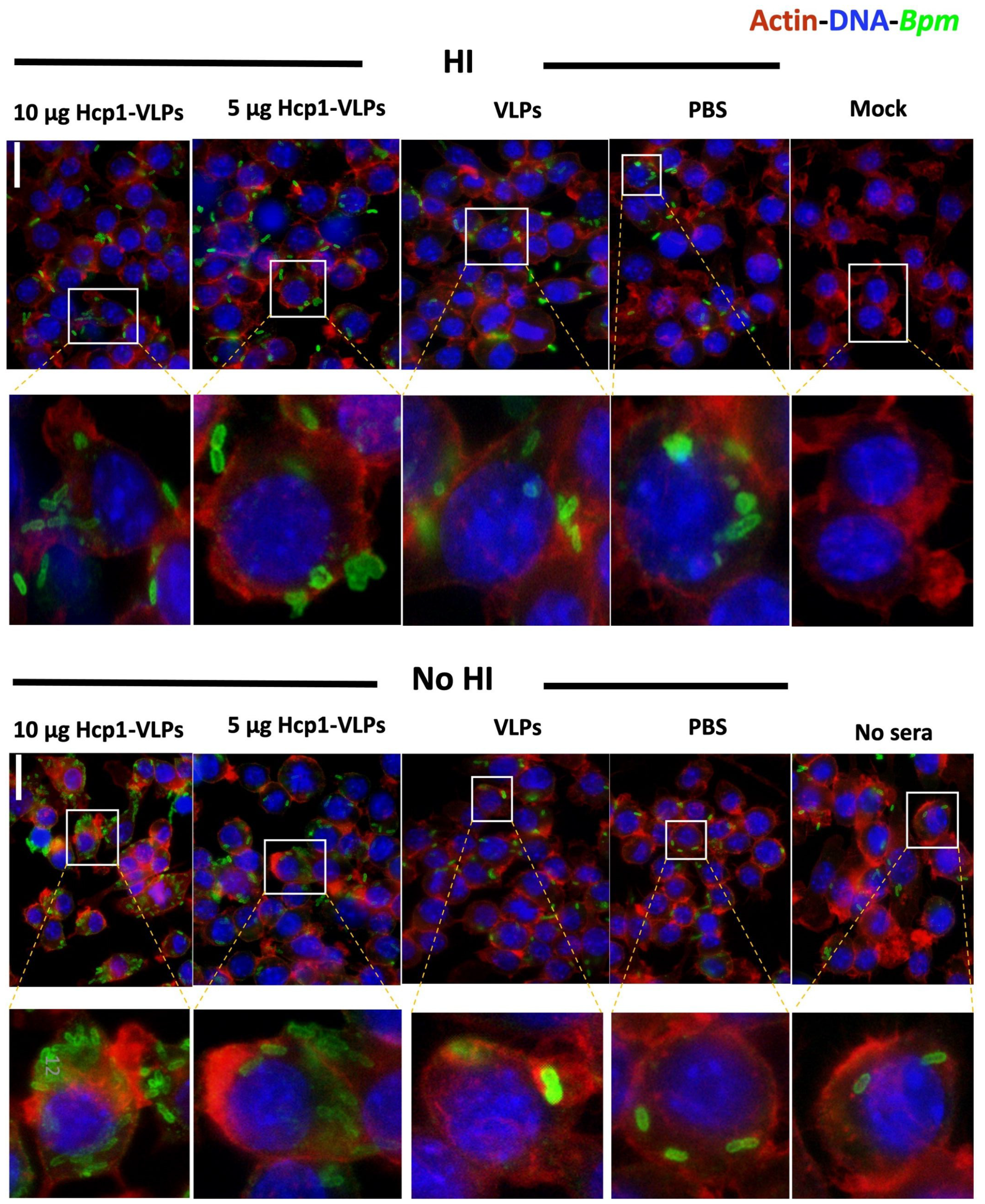
Fluorescence microscopic analysis of RAW 264.7 macrophage cells after 2 h incubation with *B. pseudomallei* K96243 in the presence of HI and non-HI immune sera from PBS, VLPs, 5 or 10 µg Hcp1-VLPs group. Cells incubated with cDMEM with (no sera) or without bacteria (mock) were used as controls. After infection, cells were fixed with paraformaldehyde, permeabilized with Triton X-100 before incubation with serum from mice vaccinated with *B. pseudomallei* PBK001. The bacterial cells, macrophage cell nuclei and actin were examined by a rabbit anti-mouse Alexa Fluor-488, DAPI and rhodamine phalloidin, respectively. Images were taken from an Olympus BX51 upright fluorescence microscope (60×) then processed using ImageJ software. The scale bar represents 20 µm.

## Discussion

VLPs represent a significant advancement in vaccine design, drug delivery and diagnostics. VLPs are currently used as alternative vaccine platforms to combat emerging infectious diseases mostly targeting viral and parasitic infections but are less frequently used against bacterial infections (28). The use of a VLP-based vaccine platform to increase the immunogenicity of heterologous antigens has been done by genetic insertion, chemical conjugation or combination of both techniques (1). For chemical conjugation, there are several methods used to display the foreign proteins and/or peptides on the VLPs surface (29). The carbodiimide chemistry is easy to operate and it is a relatively high conjugation efficient method when NHS is included (29). This protocol has been used to conjugate glucose oxidase from *Aspergillus niger* to the surface of P22-Cytochrome P450 nanoreactors for delivery of chemotherapeutic pro-drugs (30), generate meningococcal group C-tetanus toxoid vaccines (31), and conjugate the receptor-binding domain (RBD) of severe acute respiratory syndrome coronavirus 2 (SARS-CoV-2) on tobacco mosaic VLPs (32). Moreover, the EDC-NHS amino coupling is widely used to exterior decoration of drugs including folic acid and doxorubicin on cowpea mosaic virus (CPMV) for cancer therapeutic applications (33, 34).

We investigated the chemical conjugation of P22 VLPs with the known immunogenic Hcp1 protein and evaluated whether this antigen induces a potent antibody response that recognize the pathogen *B. pseudomallei*. The Hcp1 proteins were covalently conjugated to P22 VLPs using EDC-NHS amino coupling. We evaluated the conjugation efficiency by optimizing the VLPs and Hcp1 ratio as well as the protein and EDC/NHS ratio. The number of the Hcp1 molecules conjugated to the VLPs surface was counted with TEM using immunogold labeling. The optimal molar ratio of VLPs-Hcp1, EDC/NHS-Hcp1, and EDC/NHS-VLPs were 1:10, 1:100 and 1:600, respectively. The ratio of EDC/NHS to Hcp1 in the first coupling reaction and VLPs in the second coupling reaction affected conjugation efficiency. We determined that low EDC/NHS in reaction reduced conjugation efficiency due to less activated free carboxyl and amine group on Hcp1 and VLPs. In contrast, too high EDC/NHS induced self-conjugation of both proteins and can cause protein clumping. Therefore, the molar ratio of proteins and chemical needs to be optimized.

The conjugated Hcp1-VLPs formulation was evaluated for its capacity to induce antibodies with opsonizing properties. Our data demonstrated high Hcp1 specific IgG and IgA antibodies in serum of mice immunized with Hcp1-VLPs. Comparable results have been reported in other different *Burkholderia* vaccine platforms that used Hcp1 protein as antigenic component to develop *Burkholderia* vaccine candidates. For example, mice receiving three doses of gold nanoparticles conjugated Hcp1-LPS or a combination of other proteins (OmpW, OpcP, OpcP1, FlgL, and hemagglutinin) generated high titer (10^5^-10^6^ endpoint titer) of Hcp1 specific IgG (22, 24). A subunit vaccine made by conjugating Hcp1 to *Burkholderia* capsular polysaccharide (CPS) demonstrated the high serum Hcp1 specific IgG antibody titer (∼10^6^ endpoint titer) in vaccinated mice after a regimen of 3 doses (25). In addition, sera from mice immunized with Hcp1 incorporated into staphylococcal membrane vesicle plus adjuvant induced significant specific IgG antibody titer (> 51,200) (26).

Serum from Hcp1-VLP-immunized mice was only capable of promoting opsonophagocytosis when complement was present. Furthermore, baseline serum with functional complement did not promote opsonophagocytosis. These results indicate that the classical complement cascade, in which antigen-specific antibodies promote the activation of complement, it is responsible for the increase in opsonophagocytsosis observed from the immunization serum. The previous study showed similar finding that antibody-mediated enhanced uptake of *B. pseudomallei* was largely through complement activation/deposition (35). It has been previously reported that sera from Hcp1-CPS glycoconjugate vaccination promotes opsonophagocytosis (25) and the convalescent melioidosis patient sera promote antibody-dependent bacterial uptake and bacterial clearance (36). Although we were unable to evaluate our vaccine formulation in a murine model of melioidosis disease, immunological evidence suggest that Hcp1-VLPs might be protective against infection. As such, other studies using mouse models of infection also demonstrated that antibodies play a major role in protection against lethal melioidosis (27, 37, 38).

## Conclusions

Hcp1 chemically conjugated of to P22 VLPs using the EDC/NHS coupling yielded a stable nanoparticle construct for use in vaccination. This decorated VLPs-based vaccine platform was capable of inducing antibody responses and enhanced opsonophagocytosis against *B. pseudomallei* infection. This initial proof-of-concept study results provided evidence that using VLPs as a nanovaccine to delivery immunogenic proteins is advantageous and the efficiency of conjugation can be improved by optimization of VLPs, protein concentration and chemical ratio. In addition, conjugation of VLPs with multiple immunogenic proteins can enhance the protective immune responses against infectious diseases.

## Acknowledgments

This manuscript was partially supported by NIH NIAID grant AI12660101 and institutional funds from UTMB awarded to AGT. NB-T is a Pew Latin American Fellow in Biomedical Sciences. AH-S acknowledges financial support from the UNAM DGAPA-PASPA program for his sabbatical stay fellowship. The contents are solely the responsibility of the authors and do not necessarily represent the official views of the NIAID or NIH.

